# Can motional dynamics account for the cytotoxicity of beta amyloid oligomers?

**DOI:** 10.1101/2023.05.04.539338

**Authors:** Chen-Tsen Yeh, Han-Wen Chang, Wen-Hsin Hsu, Shing-Jong Huang, Meng-Hsin Wu, Ling-Hsien Tu, Ming-Che Lee, Jerry Chun Chung Chan

**Affiliations:** Department of Chemistry, National Taiwan University, No. 1, Section 4, Roosevelt Road, Taipei, 10617, Taiwan; Instrumentation Center, National Taiwan University, No. 1, Section 4, Roosevelt Road, Taipei, 10617, Taiwan; Department of Chemistry, National Taiwan Normal University, No. 88, Section 4, Ting-Chow Road, Taipei, 11677, Taiwan

**Keywords:** Aβ oligomers, reverse micelles, solid state NMR, motional dynamics

## Abstract

The underlying biophysical principle governing the cytotoxicity of the oligomeric aggregates of β-amyloid (Aβ) peptides has long been an enigma. Here we show that the size of Aβ_40_ oligomers can be actively controlled by incubating the peptides in reverse micelles. Our approach allowed for the first time a detailed comparison of the structures and dynamics of two Aβ_40_ oligomers of different size, viz., 10 and 23 nm, by solid-state NMR. From the chemical shift data, we infer that the conformation of the residues from K16 to K28 are different between the 10-nm and 23-nm oligomers. We find that the 10-nm oligomers are more cytotoxic, and the molecular motions of their charged residues are more dynamic. Interestingly, the residue A21 exhibits an unusually high structural rigidity. Our data raise the interesting possibility that the cytotoxicity of Aβ_40_ oligomers could also be correlated to the motional dynamics of the charged residues.

## Introduction

Accumulation of β-amyloid (Aβ) aggregates is a pathological hallmark of Alzheimer’s disease (AD). Soluble aggregates of Aβ have emerged as important therapeutic targets in the field of AD, owing to their neurotoxicity and ability to impair synaptic function.^[1–5]^ Many oligomeric forms of Aβ have been identified both in vitro and in vivo, ranging from dimers to large protofibrils, which contribute in different extent to neurodegeneration and synapse dysfunction.^[6–11]^ Although previous attempts to develop neutralizing antibodies for Aβ have been largely unsuccessful,^[12–15]^ Lecanemab (also known as BAN2401), which is a monoclonal antibody that targets high molecular weight (high-MW) Aβ oligomers (300–5000 kDa), has demonstrated very encouraging results in early clinical trials.^[16,17]^ Thus, high-MW AβOs are promising therapeutic targets for AD treatment.

There is a large variety of high-MW Aβ oligomers (AβOs) such as I_β_,^[18]^ amylospheroids (ASPDs),^[6]^ protofibrils,^[19,20]^ annular protofibrils,^[21]^ prefibrillar, and fibrillar oligomers.^[22]^ The size of AβOs is highly correlated with their toxicity.^[23]^ In particular, ASPDs exhibit higher neurotoxicity than both Aβ monomers and fibrils.^[6]^ It has been suggested that synthetic ASPDs shared the same mechanism of neurotoxicity as in vivo ASPDs,^[24]^ and that ASPDs have a distinct surface structure to target the presynaptic sites of neuronal cells rather than the postsynaptic sites targeted by other low-MW Aβ oligomers such as amyloid-beta derived diffusible ligands (ADDLs).^[25]^ Usually AβOs have a large distribution in size, which is accompanied with a substantial structural heterogeneity. It is very challenging to prepare structurally homogeneous AβO because of its strong propensity to further aggregate to form fibrils.^[26]^ Thus, the transient nature of AβO structure renders it very difficult to compare the biochemical results of the oligomeric aggregates of Aβ in the literature.^[27]^ Consequently, despite there are numerous oligomeric aggregates of Aβ either extracted from brain tissues or synthesized in vitro, the mechanism underlying their cell toxicity is unclear and their structure-toxicity relationship remains an enigma.^[1,6,21,28,29]^

High-MW AβOs are usually obtained by incubating the Aβ monomers at low temperature,^[11,30,31]^ or by utilizing antibodies,^[32]^ affibodies,^[33]^ other small molecules,^[34]^ and Zn^2+^ to stabilize the AβOs.^[35]^ All these methods can be categorized as the passive approach, where the size distribution of the AβOs is largely determined by the aggregation kinetics. AβOs of narrow size distribution can only be obtained by multiple rounds of filtration. Recently, we used reverse micelle (RM) to prepare the oligomeric aggregates of Aβ which are uniform in size, viz., 23 nm, and structurally monomorphic.^[36,37]^ The size of AβOs was confined by the physically dimension of the RMs. Because ASPDs derived from the brain tissues of AD patients are 4 to 18 nm in size, for which the spheres with 10–15 nm in size correlated the best with cytotoxicity,^[6]^ it is highly desirable to prepare AβOs in the regime of 10 nm. It is well known that the size of RMs can be controlled by a careful optimization of the molar ratio of the organic phase and surfactant.^[38–41]^ In this work, we were able to prepare AβOs with a diameter of 10 nm using RMs. On the basis of the results of thioflavin T (ThT) and cell toxicity assays, we showed that RM10-Aβ_40_ exhibited a distinct aggregation kinetics and it demonstrated a higher toxicity than the AβOs with a diameter of 23 nm. Furthermore, solid-state NMR results indicate that the cell toxicity may be correlated with the motional dynamics of the oligomeric aggregates of Aβ.

## Results and Discussion

### Size Control of Oligomeric Aggregates of Aβ

We previously utilized a ternary system comprising the non-ionic surfactants poly(oxyethylene) nonylphenyl ether (Igepal CO520), cyclohexane, and NH_4_OAc_(aq)_ to prepare Aβ_40_Os of 23 nm in size, viz., RM23-Aβ_40_.^[37]^ In this study, we used a 1:1 (w:w) mixture of n-hexane and cyclohexane as the organic phase for the preparation of RMs,^[40]^ whose chemical composition was optimized with respect to the CO520 concentration and the molar ratio of water to CO520, viz., w_0_ (Table S1). The dynamic light scattering (DLS) measurements revealed that the hydrodynamic diameter of the resulting RM was 10 nm **(Figure 1a)**, and the system remained stable for at least 7 days **(Figure 1b)**. Following the procedure reported previously,^[37]^ the powder sample of the Aβ_40_Os back extracted from the RM solution is henceforth referred to as RM10-Aβ_40_. We note in passing that the size of the RM can be varied in situ by adjusting the surfactant concentration and the w_0_ ratio (**Figure S1)**. From the transmission electron microscopy (TEM) images, the size distribution of RM10-Aβ_40_ was found to be 8.7 ± 2.8 nm **(Figures 1c** and **1d)**. As shown in **Figure S2**, the back extraction efficiency of the RM10-Aβ_40_ was ca. 20%.

**Figure 1.**
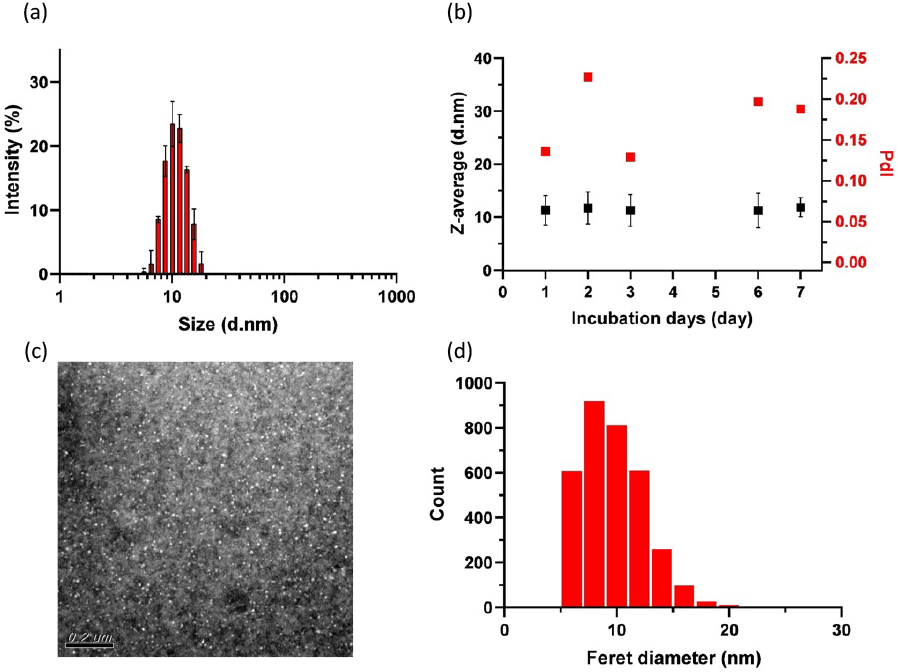
(a) DLS results of RM10-Aβ_40_. (b) Day trace of DLS data of Aβ peptides incubated in RMs. (c) TEM image of RM10-Aβ_40_ after back extraction from RMs. (d) Size distribution of the spheroids of RM10-Aβ_40_ deduced from the TEM images.

### Self-aggregation and Toxicity of RM10-Aβ_40_

RM10-Aβ_40_ formed fibrils after resuspension in phosphate buffer for 60 hours (**Figure S3**). The same result was observed for RM23-Aβ_40_.^[37]^ As shown by the ThT excitation profile, the spheroids of RM10-Aβ_40_ were ThT positive, showing an absorption maximum in the region of 440–450 nm (**Figure 2a**). Their aggregation kinetics were traced by monitoring the ThT emission at 490 nm. The results of RM23-Aβ_40_ did not have any significant variation in the ThT fluorescence intensity even after fibril formation (**Figure S4a**). This observation revealed that the β-sheet structure of the RM23-Aβ_40_ spheroids were comparable to that of the subsequently formed fibrils. By contrast, the ThT results of RM10-Aβ_40_ exhibited a distinct lag phase, whose duration was found to be dose dependent (**Figure S4b**). After analyzing the kinetic profiles by AmyloFit,^[42]^ we found that the results of the three lowest concentrations (×0.1, ×0.25, and ×0.5) could be reasonably fitted by the multi-step secondary nucleation model using a single set of rate constants for global fitting (**Figure 2b**).^[43]^ As the concentration increased to ×0.75, considerable deviation from the model was observed (**Figure S4c**). At the highest concentration in this study, viz., ×1, the profile was more consistent with the saturation elongation model (**Figure S4d**).^[44]^ Apparently, the β-sheet structure of RM10-Aβ_40_ was less ordered than that of RM23-Aβ_40_. During the lag phase, the spheroids of RM10-Aβ_40_ might undergo either the dissolution process to release Aβ_40_ monomers, or the coalescence process leading to fibril formation. As the concentration of RM10-Aβ_40_ increased to ×1, the latter process became kinetically more favorable. The coalescence process of the RM10-Aβ_40_ spheroids can be aptly described by the saturation elongation model originally derived for Aβ_40_ monomers, where the spheroids were the basic strucutral unit instead of the monomers. However, it is not trivial to interpret the numerical values of the rate constants extracted for the models.

**Figure 2.**
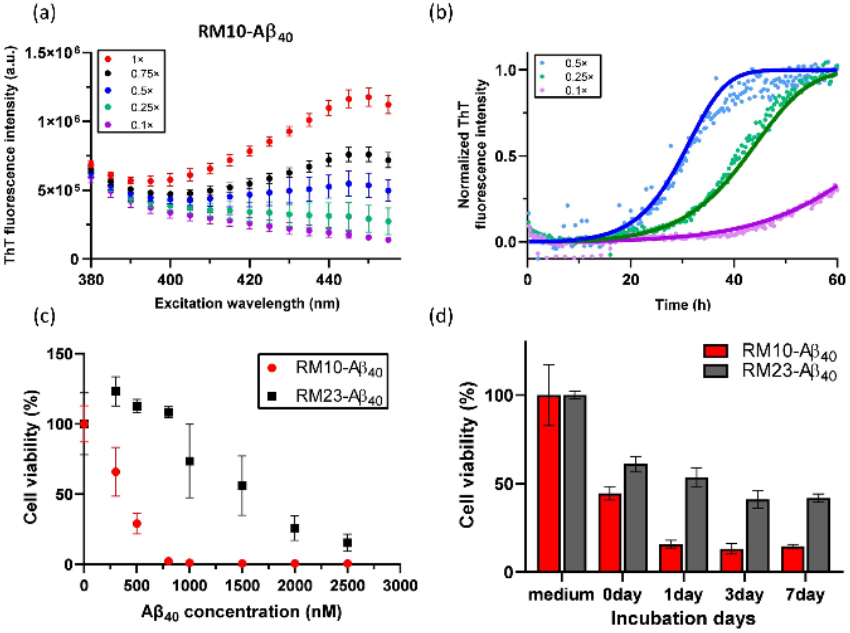
(**a**) Excitation profile of ThT fluorescence for different concentrations of RM10-Aβ_40_. (**b**) Normalized ThT data at different concentrations of RM10-Aβ_40_. The solid curves were calculated based on the multi-step secondary nucleation model using a single set of rate constants for global fitting. (**c**) Cytotoxicity of RM10-Aβ_40_ and RM23-Aβ_40_ assessed by alamarBlue assay. Data were the mean of ten replicates for each concentration relative to the control (DMEM only). Error bars indicate standard deviation of the mean cell viability values. (**d**) Cytotoxicity of RM10-Aβ_40_ and RM23-Aβ_40_ with different incubation periods in RMs.

Prior research has reported that there is a correlation between the size of AβOs and their cytotoxicity.^[23]^ To evaluate the cytotoxicity of RM10-Aβ_40_ and RM23-Aβ_40_, we performed the alamarBlue assay on 293T cell lines for a series of samples diluted against 10 µM of CO520, which is the typical residual concentration of CO520 among all samples. We confirmed that 10 µM of CO520 did not exhibit any significant cell cytotoxicity (**Figure S5**). As shown in **Figure 2c**, RM10-Aβ_40_ was more toxic than RM23-Aβ_40_, which is consistent with the literature results.^[23]^ We incubated Aβ_40_ peptides in RMs for 0 to 7 days, and observed the cytotoxicity of the RM10-Aβ_40_ and RM23-Aβ_40_ hence extracted for each of the incubation period (**Figure 2d**). As the incubation time increased, both RM10-Aβ_40_ and RM23-Aβ_40_ developed cytotoxicity. Again, RM10-Aβ_40_ showed significantly stronger cytotoxicity than RM23-Aβ_40_. The cytotoxicity of RM10-Aβ_40_ reached its plateau value earlier than RM23-Aβ_40_, presumably because the smaller water droplets in RM10-Aβ_40_ would lead to faster aggregation kinetics. Nonetheless, the cytotoxicity of both the samples reached their plateau values after 7 days. This observation justified our protocol to incubate the Aβ peptides in RMs for 7 days. It is well documented the cytotoxicity of Aβ is in the order of low-MW AβOs > high-MW AβOs > Aβ fibrils ≫ Aβ monomers. It has long been an enigma why the cytotoxicity of AβOs decreased as its size increased. Therefore, it would be very interesting to probe whether there is any significant structural variation between RM10-Aβ_40_ and RM23-Aβ_40_.

### Comparison of the ^13^C Chemical Shifts of RM-Aβ_40_

Four selectively ^13^C enriched RM10-Aβ_40_ samples with an enrichment level of 60% were synthesized for solid-state NMR study. A typical ^13^C–^13^C correlation spectrum acquired for RM10-Aβ_40_ incubated for 7 days is shown in **Figure 3a**. Although RM10-Aβ_40_ are transient species, the spectra of RM10-Aβ_40_ is dominated by a single set of cross peaks (**Figures S6–S9**), indicating that the oligomers with a size of 10 nm are still structurally monomorphic. This observation supports the conjecture that the Aβ_40_ peptides aggregate via a single pathway during the nucleation process in RMs. The ^13^C NMR spectra of RM10-Aβ_40_ and RM23-Aβ_40_ are rather similar in line widths (**Figures 3b, S10**). Although the chemical shift data of E11-C^β^, E11-C_α_, L17-C_β_, E22-CO, D23-CO, N27-CO, N27-C_α_, N27-C_β_, K28-C^β^, and I31-C^β^ were not determined for RM10-Aβ_40_ due to insufficient signal intensity and/or spectral resolution, the secondary chemical shift analysis of the other observed signals indicated that RM10-Aβ_40_ also adopted the structural motif of β_1_-turn-β_2_ (**Figure S11**), which is the same as that of RM23-Aβ_40_ and Aβ_40_ fibril structures.^[37]^ Our analysis revealed that the backbone ^13^C chemical shift difference between RM10-Aβ_40_ and RM23-Aβ_40_ was relatively minor in the β_2_ region (**Figure 3c**). For RM10-Aβ_40_, the poor spectral resolution of the N27 signals rendered the comparison of its chemical shifts with those of RM23-Aβ_40_ very difficult. It is rather surprising that the structural order at D23 was somewhat higher for RM10-Aβ_40_ than RM23-Aβ_40_. That is, there was only a single set of cross-peak signals at D23 observed for RM10-Aβ_40_, whereas multiple peaks were observed for RM23-Aβ_40_ (**Figure S12a**). Interestingly, the chemical shifts of D23 determined for RM10-Aβ_40_ were different from those reported for other Aβ_40_ aggregates (**Figure S12b**). Altogether, the chemical shift difference between RM10-Aβ_40_ and RM23-Aβ_40_ was very substantial from D23 to N27, which is a strong indication that the backbone conformation at the turn region was altered significantly as the size of AβOs increased from 10 to 23 nm. In the β_1_ region, the chemical shifts of K16 also exhibited a large deviation between RM10-Aβ_40_ and RM23-Aβ_40_.

**Figure 3.**
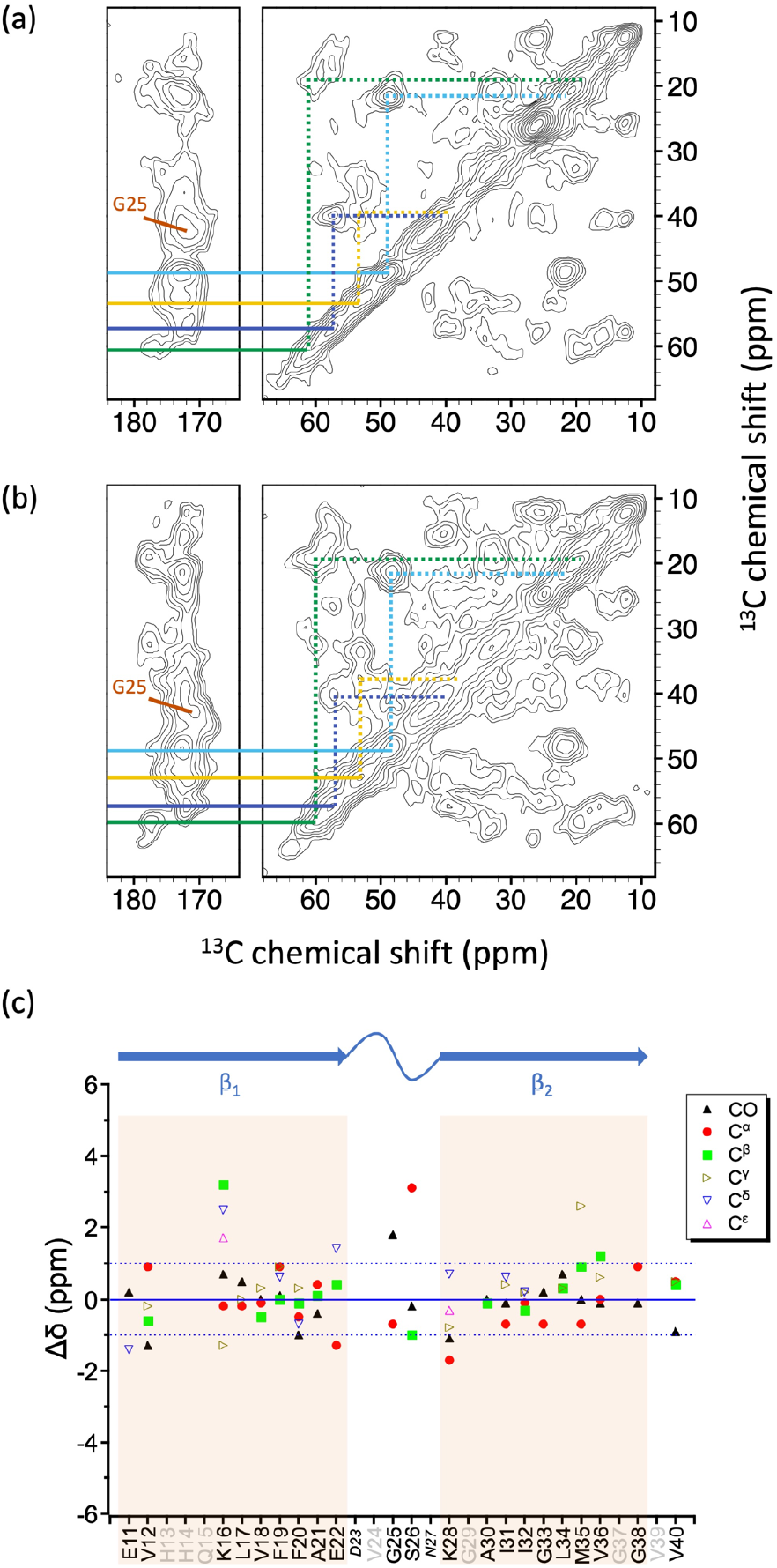
Comparison of typical NMR spectra for (**a**) RM10-Aβ_40_ (this work) and (**b**) RM23-Aβ_40_ (adapted from our previous work).^[37]^ Colored lines denote signal assignments for K16 (yellow), A21 (cyan), G25 (brown), I32 (blue), and V40 (green). The contour levels were increased by a factor of 1.4 successively, where the base levels were set to 4× root-mean-square noise. The spectra show no significant different in terms of pattern and line width. (**c**) Difference of chemical shift (Δ!) between RM10-Aβ_40_ and RM23-Aβ_40_. Residues not ^13^C enriched are shown in grey. The signals of the italicized residues D23 and N27 were unassigned in the spectrum of RM23-Aβ_40_ and RM10-Aβ_40_, respectively, due to poor spectral resolution. The β-sheet regions, defined based on the chemical shift data from RM23-Aβ_40_, are highlighted in light orange.

### Variation in ^13^C Signal Intensities of RM-Aβ_40_

All the 2D homonuclear ^13^C correlation spectra acquired for RM-Aβ_40_ were based on the techniques of ^13^C{^1^H} cross polarization (CP) and dipolar-assisted rotational resonance (DARR). It is well known that their signal intensities are highly sensitive to the motional dynamics of the spin system. Thus, the intensities of the cross peaks should reveal the difference in motional dynamics between RM10-Aβ_40_ and RM23-Aβ_40_. We rescaled the cross peak intensities of the RM-Aβ_40_ spectra using their cross peak intensities of the backbone carbons of I32, G33, V34, and M35, where the signals of the β_2_ region were well resolved in the RM10-Aβ_40_ and RM23-Aβ_40_ spectra (**Figures S13–S14)**. For convenience, we summarize the normalized cross-peak intensities with the most significant variation in **Figure 4**. Accordingly, RM23-Aβ_40_ has more observable cross-peak signals than RM10-Aβ_40_ for the sidechains of K16, E22, K28, I31, and for the backbone of V40. The extent of the cross-peak attenuation was most prominent for K16 and A21 for RM10-Aβ_40_. It is likely that the side-chain molecular motions at K16 was more dynamic for RM10-Aβ_40_, although it could partly be the consequence of the low signal-to-noise ratio. The cross peaks of A21 were much more intense than other signals including those of A30. This implied that the residue A21 had the least motional dynamics for RM23-Aβ_40_. The intensities of A21 were attenuated for RM10-Aβ_40_, whereas their chemical shifts still resembled closely to those of RM23-Aβ_40_. We surmised that the molecular motions at A21 mainly manifested the motional effects of other residues in proximity. These observations suggested that the structural order of RM23-Aβ_40_ was higher than RM10-Aβ_40_, which is consistent with our ThT data. At the current stage, we are not able to construct a detailed model for the motional dynamics of RM-Aβ_40_. Nonetheless, we can safely conclude that the molecular motions associated with the sidechains of the charged residues of RM10-Aβ_40_ were more dynamic than those of RM23-Aβ_40_.

**Figure 4.**
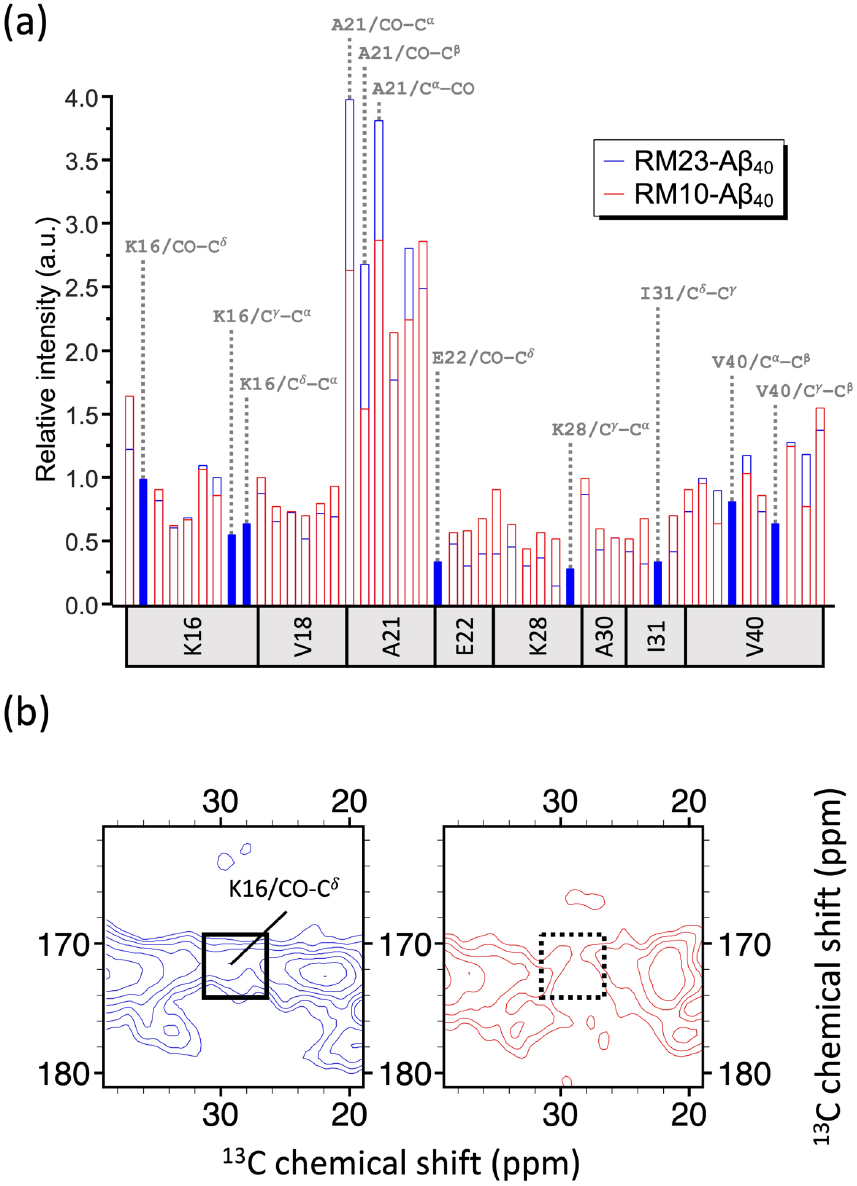
(a) Histogram of the scaled cross-peak intensities of the spectra of RM23-Aβ_40_ (blue) and RM10-Aβ_40_ (red) for selected residues. The cross-peak intensities of the backbone carbons of I32, G33, V34, and M35, located in the β2 region, were used to rescale the cross-peak intensities of other residues. Solid bars with assignments on top denote the cross peaks with significant variation. (b) Excerpt spectra highlighting the difference of the cross-peak K16/CO-C°.

### Cytotoxicity of RM10-Aβ_40_

Various mechanisms of cytotoxicity have been proposed for the oligomeric aggregates of Aβ peptides. The experimental results of oligomer-specific antibody suggested that a common backbone structure may exist for the oligomers of various amyloidogenic peptides and the neurotoxicity of AβOs is independent of the side chain conformation.^[1]^ Along this line of argument, the fibril-like β-sheet structure has been suggested to be the common structural feature of various toxic amyloid intermediates.^[18]^ The channel-like structure formed by amyloid peptides reconstituted in bilayer membrane is a well-received mechanistic rationalization for the pathophysiological and degenerative effects of amyloid peptides.^[45]^ On the other hand, it has been argued that the distinct surface tertiary structure of AβOs, which is presumably determined by the sidechain structure of the residues near the surface, is responsible for the toxic effects.^[6]^ A more recent study indicated that the cytotoxicity of AβOs is regulated by its surface structure. In particular, the solvent exposure of the hydrophobic β_1_-turn region has been considered as the key determinant of cytotoxicity.^[46]^

While the structure-toxicity relationship associated with AβOs remains actively sought after, the effects of motional dynamics on cytotoxicity have started to be aware of. It has been mentioned that AβOs are more toxic than fibril fragments because they can be endocytosed efficiently, which is attributed to the more dynamic state of oligomers than fibrils.^[47]^ The disordered N-terminus of AβOs was also found to be crucial in determining the toxicity.^[48]^

In this work, we prepared two AβOs of 10 and 23 nm in size. The former was more cytotoxic than the latter. The ThT results indicated that the β-sheet content is higher in the latter. That is, the molecular structure of the 23-nm AβOs was more fibril like. The NMR signal line widths were comparable between RM23-Aβ_40_ and RM10-Aβ_40_, implying that their structural orders were similar. Because the chemical shifts observed for the β_1_-turn region had larger deviation between RM10-Aβ_40_ and RM23-Aβ_40_, e surmised that there was a major conformational difference in the turn region as the size of AβOs increased from 10 to 23 nm. Thus, our NMR data are consistent with the notion of structure-toxicity relationship.

The intensity data had shed light on the difference in dynamic state of the AβOs. The peaks missing in the 10-nm AβOs mostly came from the sidechain carbons of the charged residues in the β_1_-turn region. It is reasonable to suggest that the motions associated with these charged residues are different for the 10- and 23-nm AβOs. As the size of AβOs increased to 23 nm, which is more fibril like, the molecular motions of those charged residues became more restricted, rendering their NMR signals observable.

Notably, the cross-peak intensities of A21 are significantly higher than all other residues for 10- and 23-nm AβOs. This observation unequivocally showed that A21 exhibited a high degree of rigidity or structural stability. One possible scenario is that A21 is a molecular “hinge”, allowing for large-scale movements of the β-sheets of the AβOs. The structural rigidity at A21 would place a strong experimental constraint for any subsequent modeling studies of the motional dynamics of AβOs.

Overall, the NMR data in this work allowed a direct comparison of the chemical states of 10- and 23-nm oligomeric aggregate of Aβ_40_. We observed that the backbone conformation was different in the β1 and turn regions and that the motional dynamics of the more toxic 10-nm AβOs were higher than the 23-nm AβOs. While we do not intend to undermine the importance of structure-toxicity relationship for AβOs, we are led to another plausible hypothesis that sidechain motions could play an important role in modulating the cytotoxicity of the oligomers. More experiments are required to verify this intriguing hypothesis.

## Conclusion

By controlling the size of RMs, we were able to actively control the size of the oligomer aggregates of Aβ peptides prepared in RMs. The AβOs of 10 nm in size was more cytotoxic than those of 23 nm. The latter was more fibril like in structure. The region of β_1_-turn exhibited a substantial variation in conformation between 10- and 23-nm AβOs. While this observation is consistent with the results of the literature, we found that the motional dynamics at the charged residues were more severe for the smaller and more toxic AβOs. In addition, the residue A21 appeared to play an important role in the collective motions of the AβOs which may be correlated with the extent of cytotoxicity.

## Acknowledgements

This work was financially supported by the Ministry of Science and Technology (MOST 108-2113-M-002-001). The NMR and TEM measurements were carried out at the Instrumentation Center of National Taiwan University, supported by the Ministry of Science and Technology. We thank Ya-Yun Yang for her help in TEM measurements.

## Figures

**Fig. S1.**
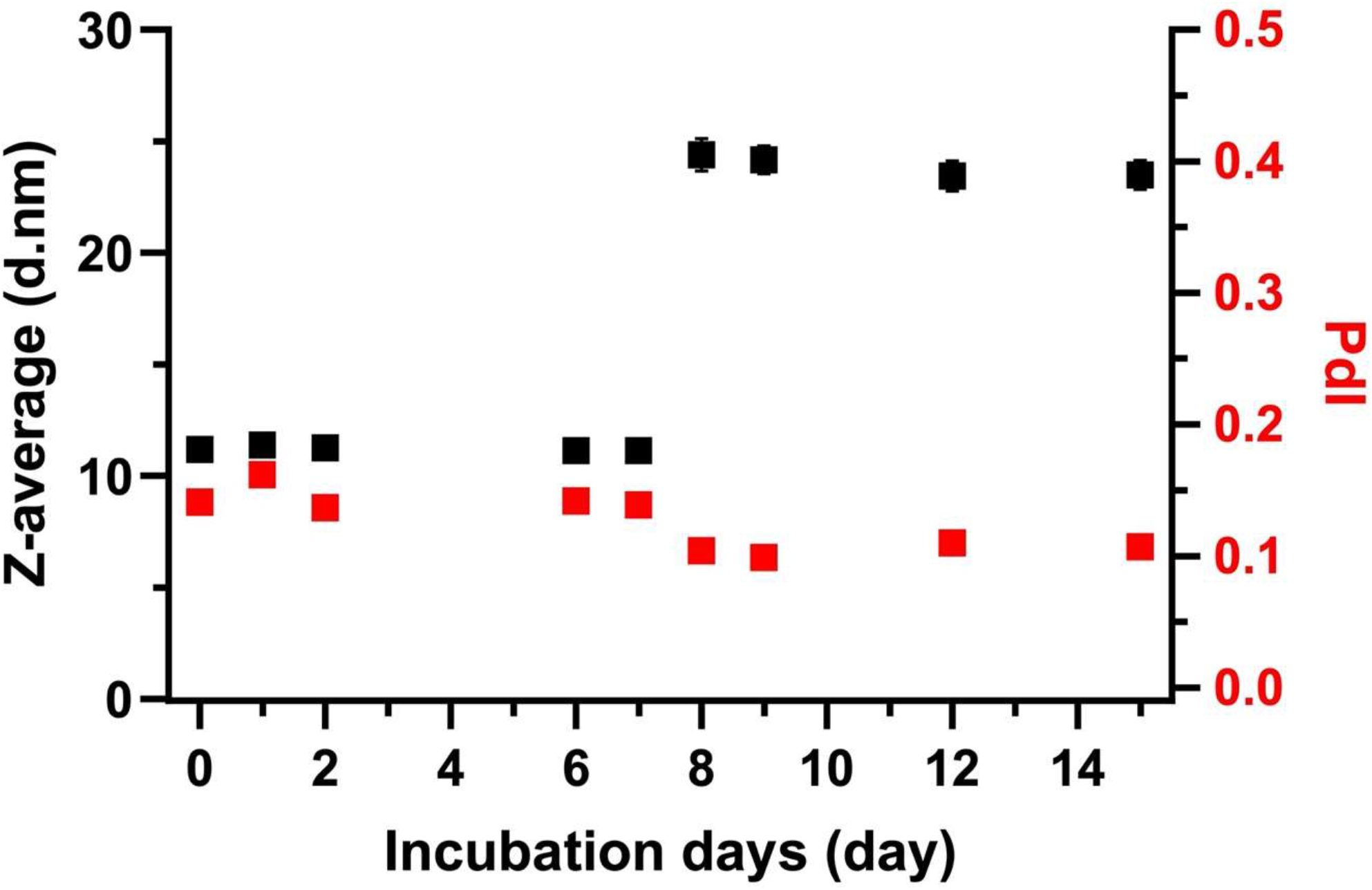
DLS data of Aβ peptides incubated in RMs. The RM solution was prepared by dissolving 1.45 g of Igepal CO520 in a mixture of 3.66 mL of *n-*hexane and 3.13 mL of cyclohexane, followed by adding 329 μL freshly prepared Aβ_40_ monomer solution (40 µM). After incubating for seven days, 4.41 mL of *n*-hexane and 3.79 mL of cyclohexane were added slowly, followed immediately by the addition of 175 uL of Aβ monomer solution (40 µM). The average size of the RMs for the first 7 days was ca. 10 nm. After increasing the amount of the organic phase and the *w*_0_ ratio, the size increased to ca. 20 nm. As expected, we could actively control the size of RMs by adjusting the chemical composition of the RM system.

**Fig. S2.**
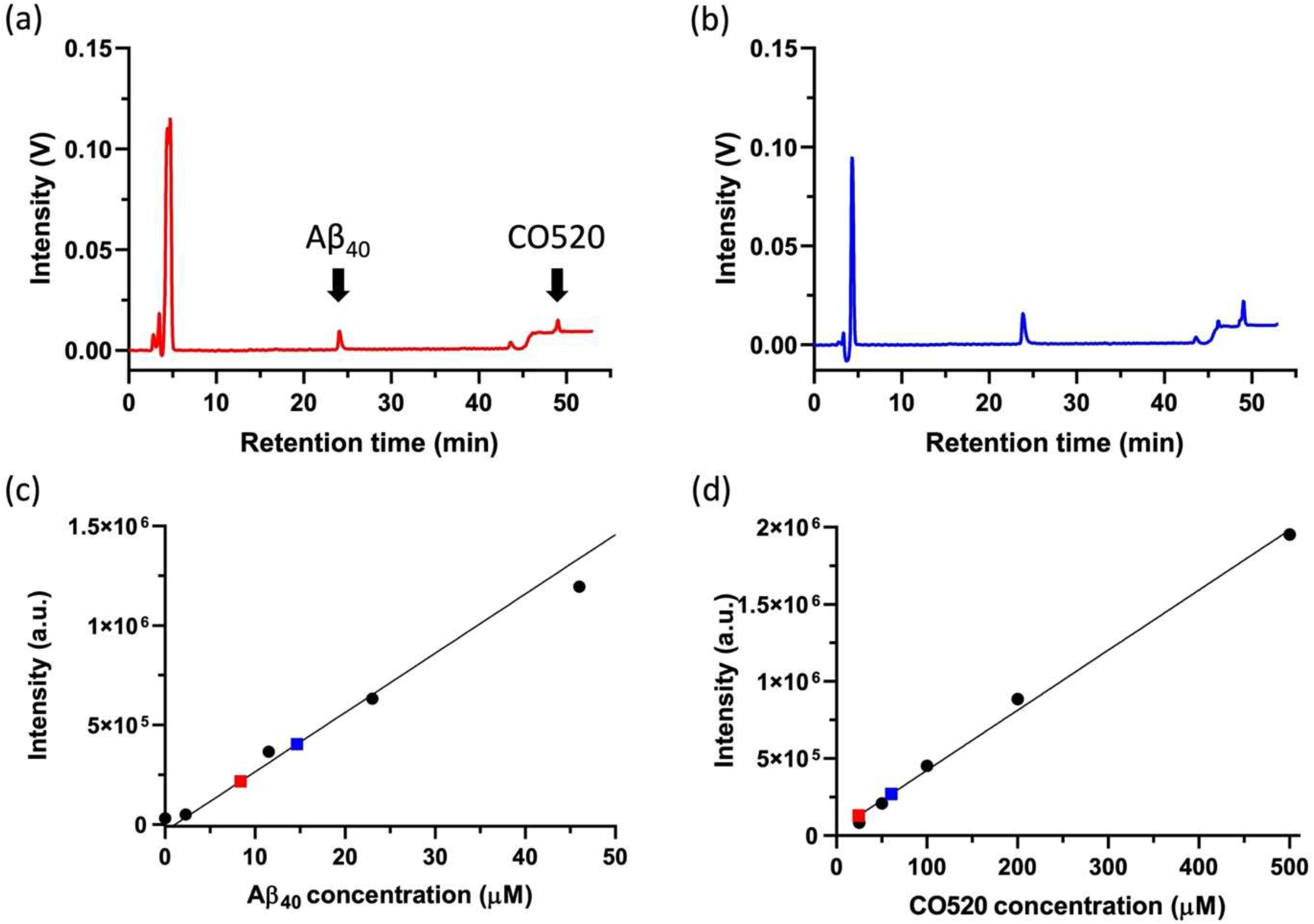
Quantification analysis of Aβ and CO520 to estimate back-extraction efficiency. The HPLC chromatograms of (**a**) RM10-Aβ_40_ and (**b**) RM23-Aβ_40_ acquired by an analytical C18 column, where the black arrows indicate the retention time of Aβ_40_ and CO-520, respectively. (**c**) Aβ_40_ signal integrals of RM10-Aβ_40_ (red square) and RM23-Aβ_40_ (blue square), together with the reference data acquired for the solutions of Aβ_40_ monomers with known concentrations (black circles). The Aβ_40_ concentrations of RM10-Aβ_40_ and RM23-Aβ_40_ were quantified as 8.4 μM and 14.7 μM, respectively. The back extraction efficiency of RM10-Aβ_40_ and RM23-Aβ_40_ were quantified as 23.2 % and 17.6 %. (d) CO520 signal integrals of RM10-Aβ_40_ (red square) and RM23-Aβ_40_ (blue square), together with the reference data acquired for the solutions of CO520 with known concentrations (black circles). The CO520 concentrations of RM10-Aβ_40_ and RM23-Aβ_40_ were quantified as 24.4 μM and 60.6 μM, respectively. Hence, the molar ratio of CO520 to Aβ_40_ ratio were calculated to be 2.9 and 4.1 for of RM10-Aβ_40_ and RM23-Aβ40, respectively. At the Aβ_40_ concentration of 2.5 μM, the concentration of CO520 was less than 10 μM for both RM10-Aβ_40_ and RM23-Aβ_40_.

**Fig. S3.**
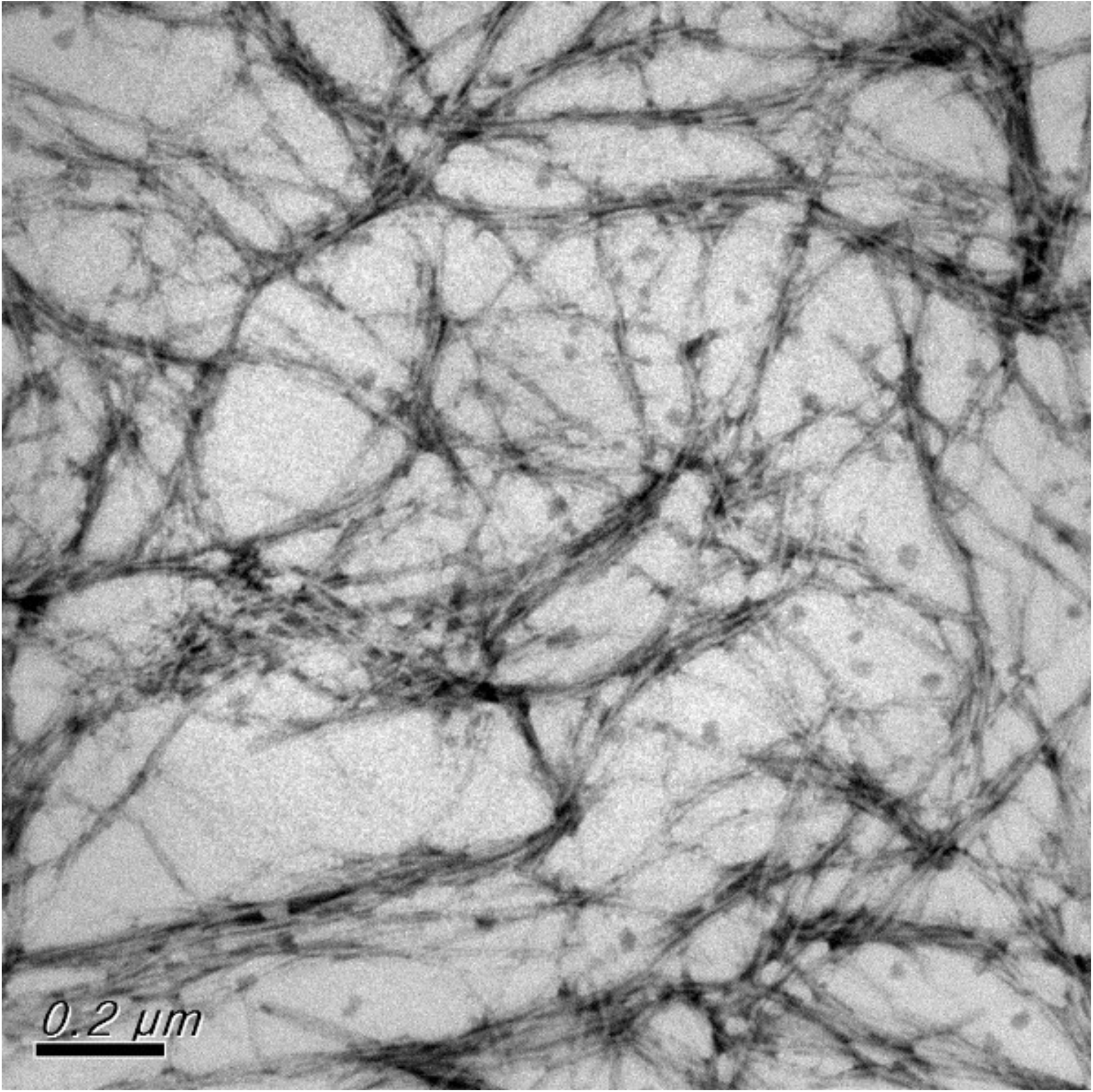
Typical TEM image of the fibrils formed through the self-aggregation of RM10-Aβ_40_, which were incubated at 37 °C and shook (15 sec every 15 min) for 60 h.

**Fig. S4.**
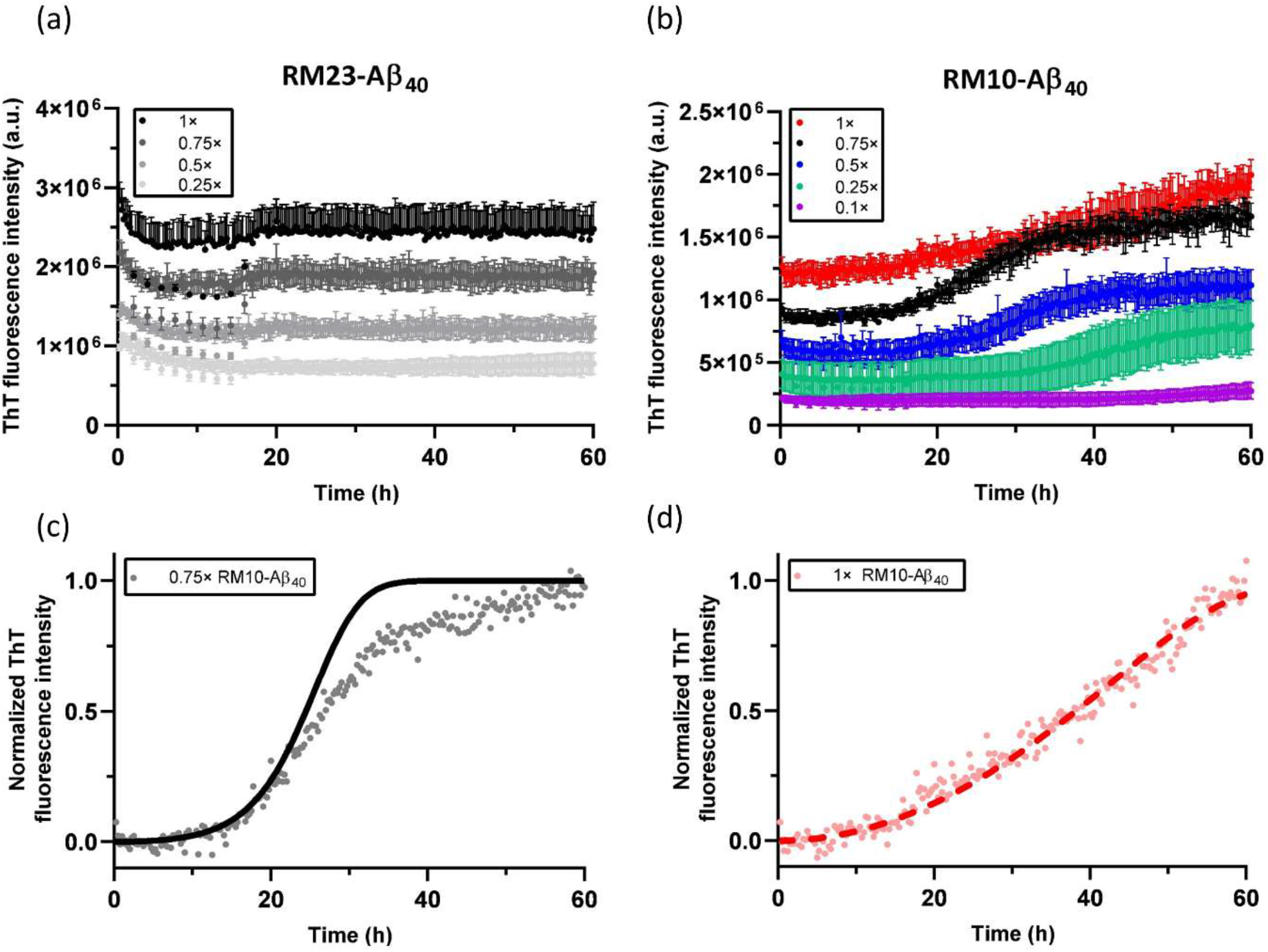
ThT profiles of (**a**) RM23-Aβ_40_ and (**b**) RM10-Aβ_40_. The nominal concentrations of RM10-Aβ40 and RM23-Aβ40 were set to be 100 µM, from which the solutions of other concentrations were prepared by dilution. The ThT data represent the mean of four replicates for each concentration. (**c**) The normalized ThT data for RM10-Aβ_40_ at a dilution factor of 0.75 demonstrated considerable deviation from the fitting curve calculated based on the global fitting parameters employed for Figure 2b of the main text. (**d**) The ThT data of the mother liquor of RM10-Aβ_40_ was normalized arbitrarily. The data could only be satisfactorily fitted by the saturating elongation model. As the concentration of RM10-Aβ_40_ varied, our data showed that there was definitely a change in the aggregation mechanism.

**Fig. S5.**
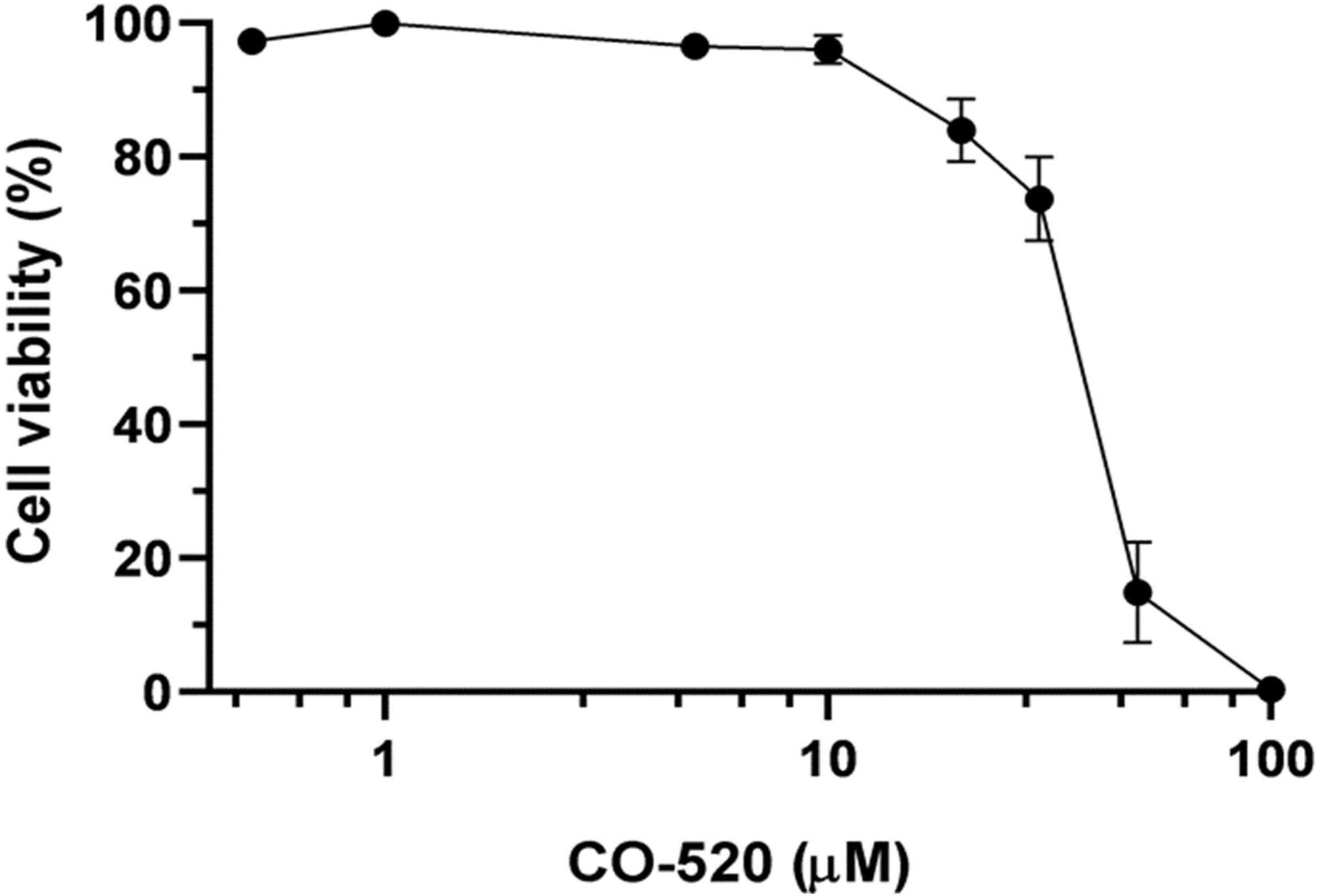
Toxicity of CO520 assessed by alamarBlue assay. Cell viability was measured as the reduction of the alamarBlue dye by viable cells after 24-hour exposure to CO520 solution at indicated concentration. The cell toxicity of CO520 at 10 µM was very minor.

**Fig. S6.**
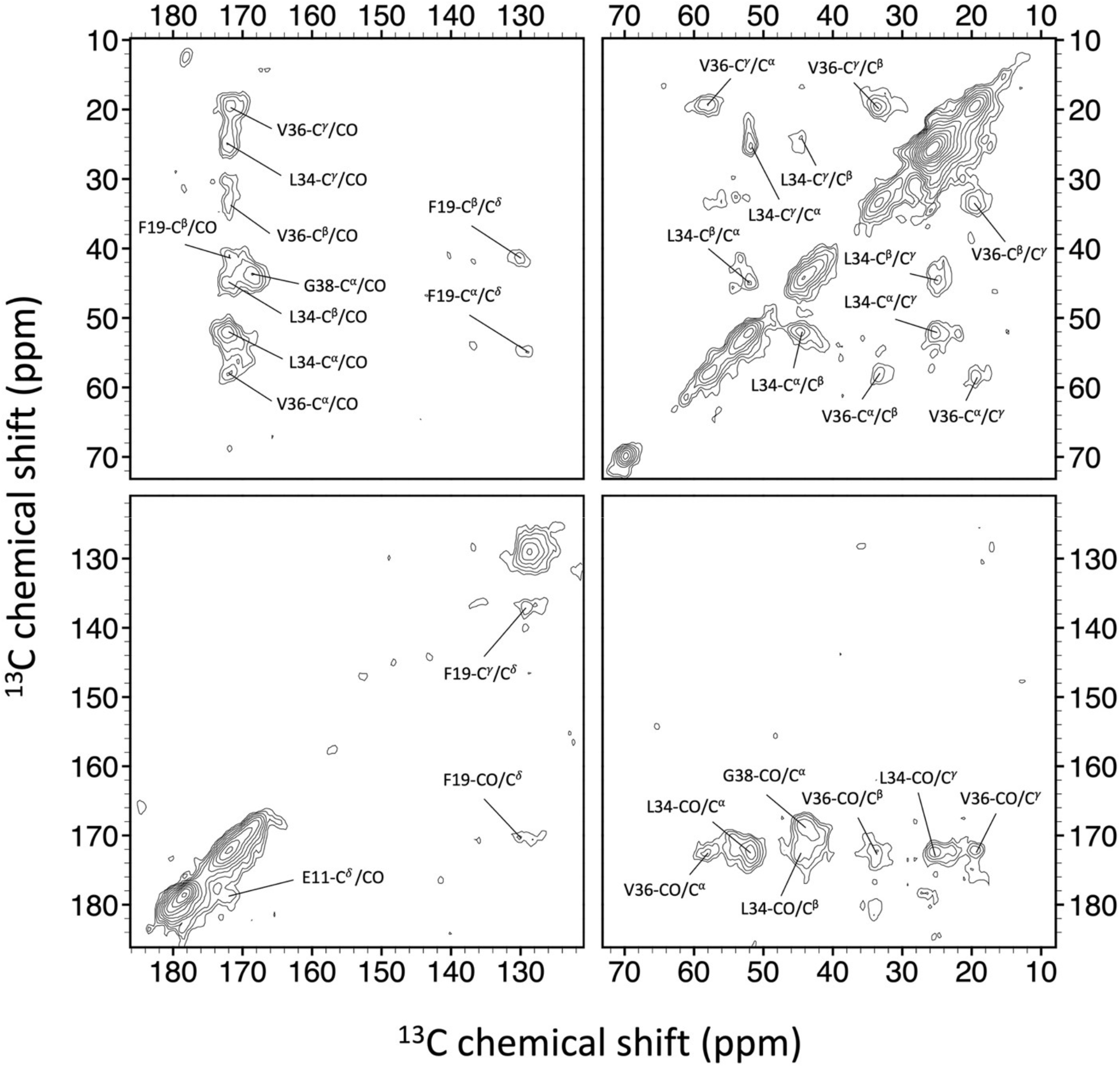
Spectrum assignment for the ^13^C–^13^C correlation spectrum of RM10-Aβ_40_ with S1 labeling scheme (see Table S2). The contour levels were increased by a factor of 1.4 successively, where the base levels were set to 4× root-mean-square noise. The processing parameters were described in Materials and Methods.

**Fig. S7.**
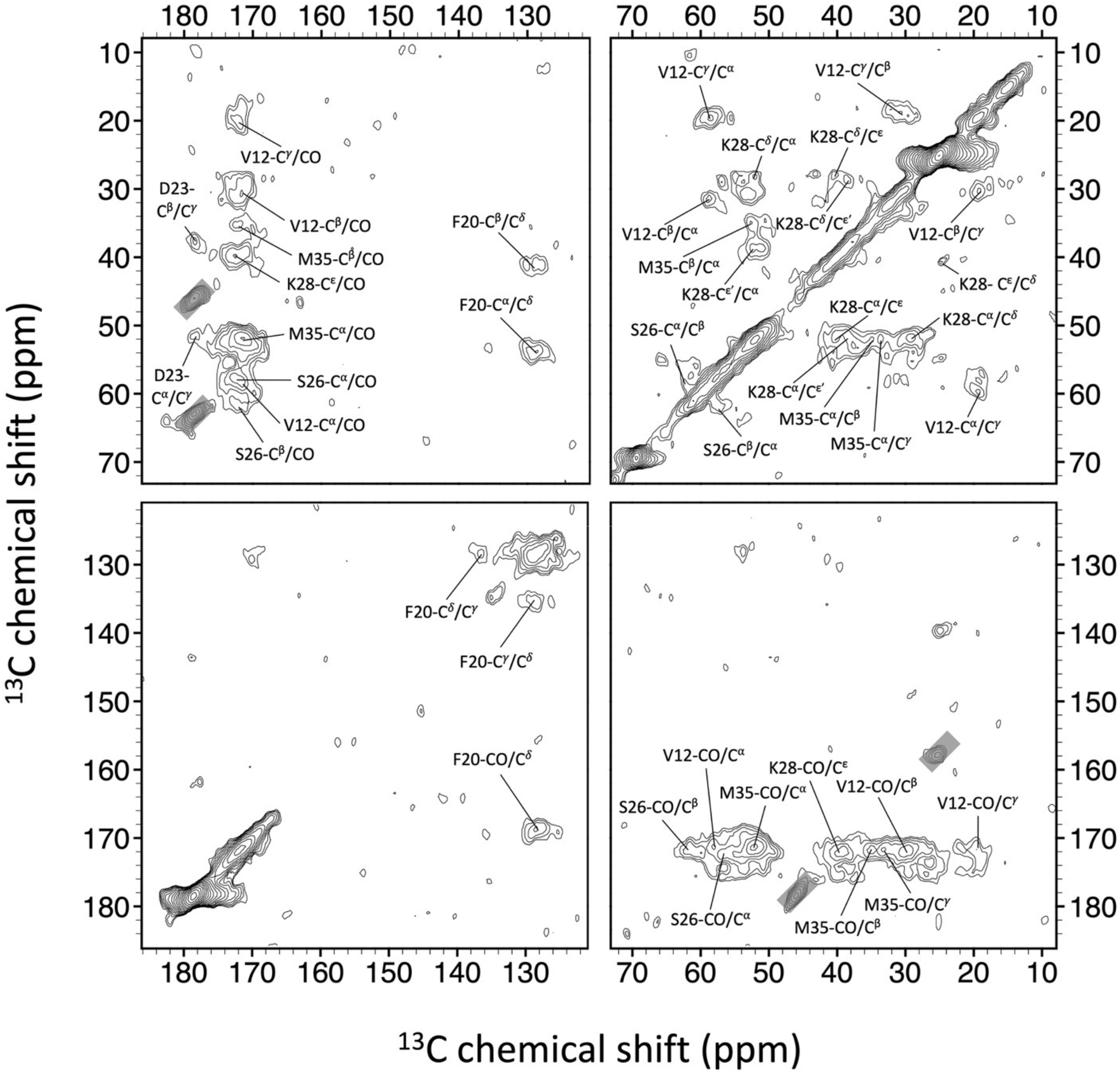
Spectrum assignment for the ^13^C–^13^C correlation spectrum of RM10-Aβ_40_ with S2 labeling scheme. The tilted rectangular boxes indicate the spinning sidebands.

**Fig. S8.**
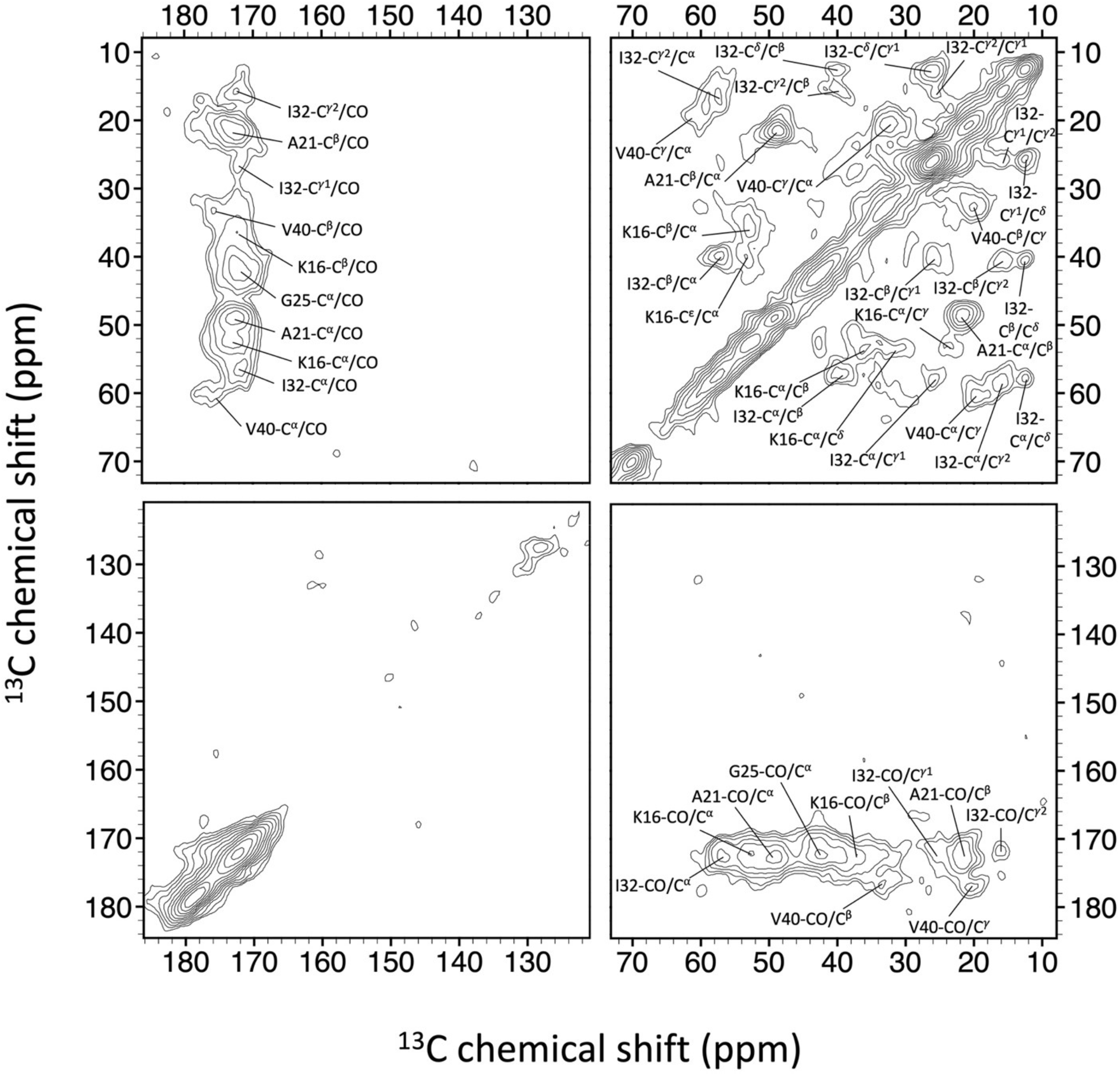
Spectrum assignment for the ^13^C–^13^C correlation spectrum of RM10-Aβ_40_ with S3 labeling scheme.

**Fig. S9.**
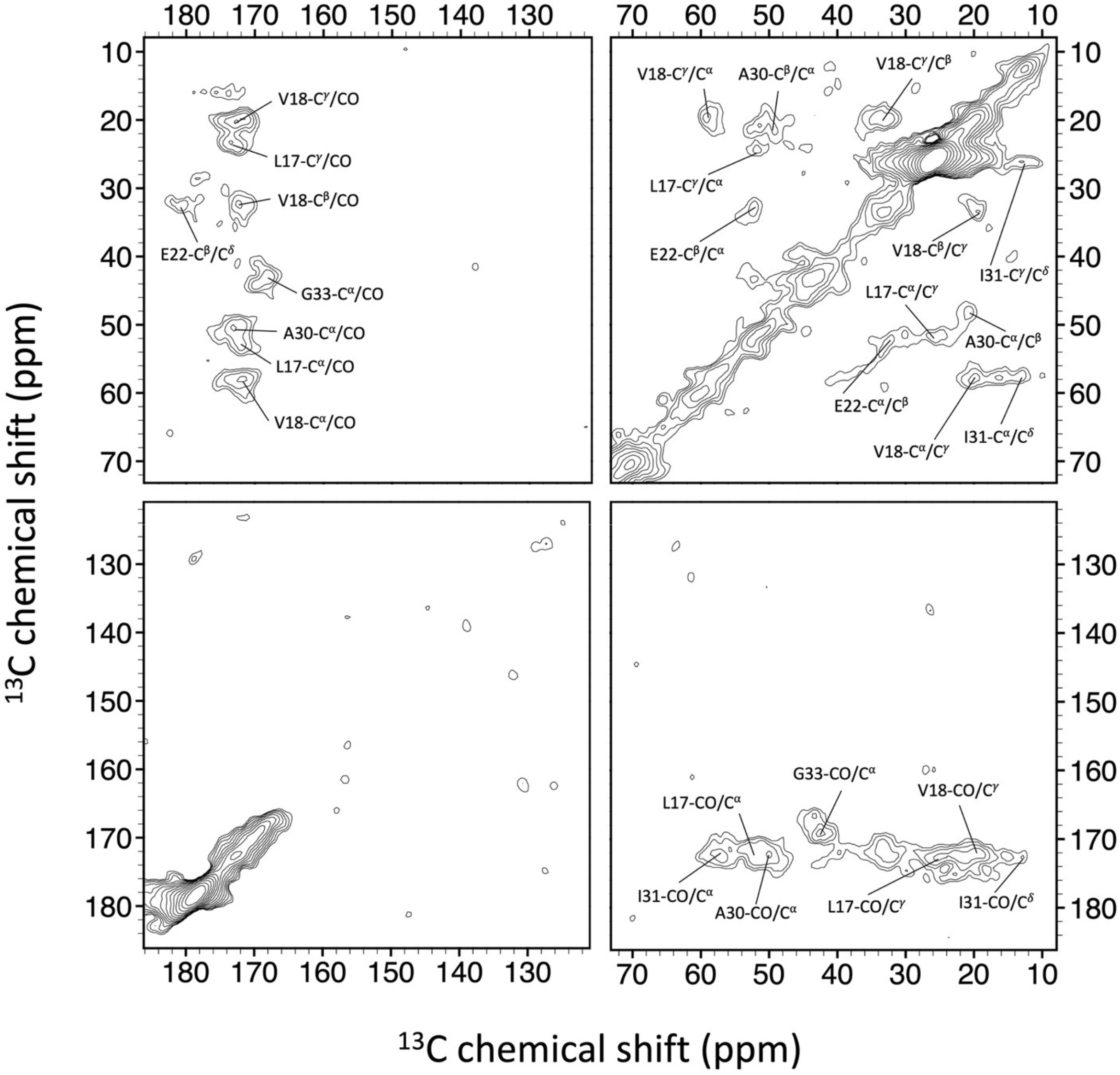
Spectrum assignment for the ^13^C–^13^C correlation spectrum of RM10-Aβ_40_ with S4 labeling scheme.

**Fig. S10.**
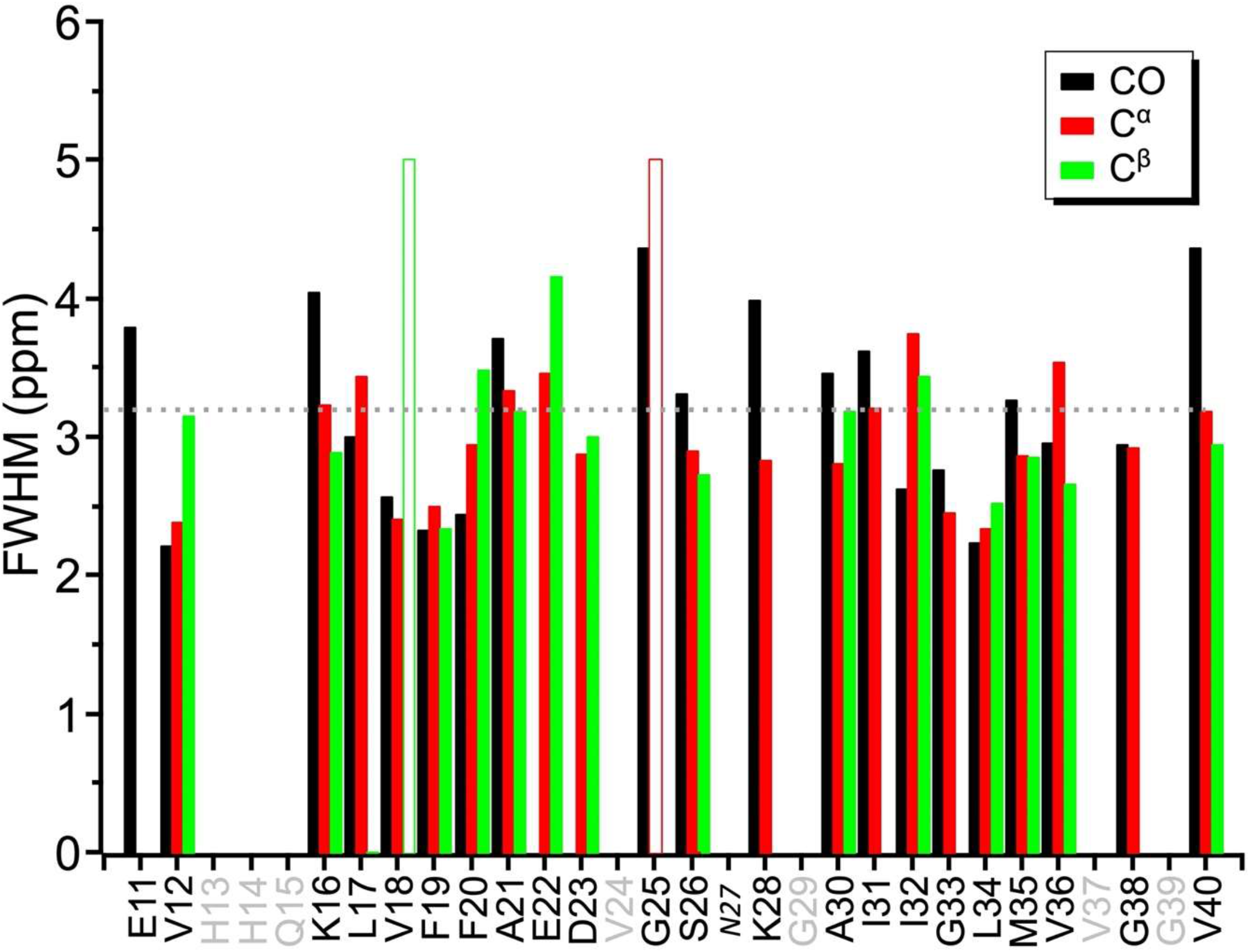
Full width at half maximum (FWHM) data for RM10-Aβ_40_. The FWHM data of E11-C_α_, E11-C^β^, L17-C^β^, F19-C_α_, F19-C^β^, E22-CO, D23-CO, N27, K28-C^β^, I31-C^β^ were not determined due to the limited spectral resolution. The dashed lines denote the average FWHM of 3.2 ppm. The open bars denote FWHM > 5 ppm, which were not included in calculating the average FWHM. Residues not ^13^C enriched are shown in grey. The signals of the italicized residues (N27) were unassigned in the spectrum of RM10-Aβ_40_ due to poor resolution and the absence of any prominent relay cross peaks.

**Fig. S11.**
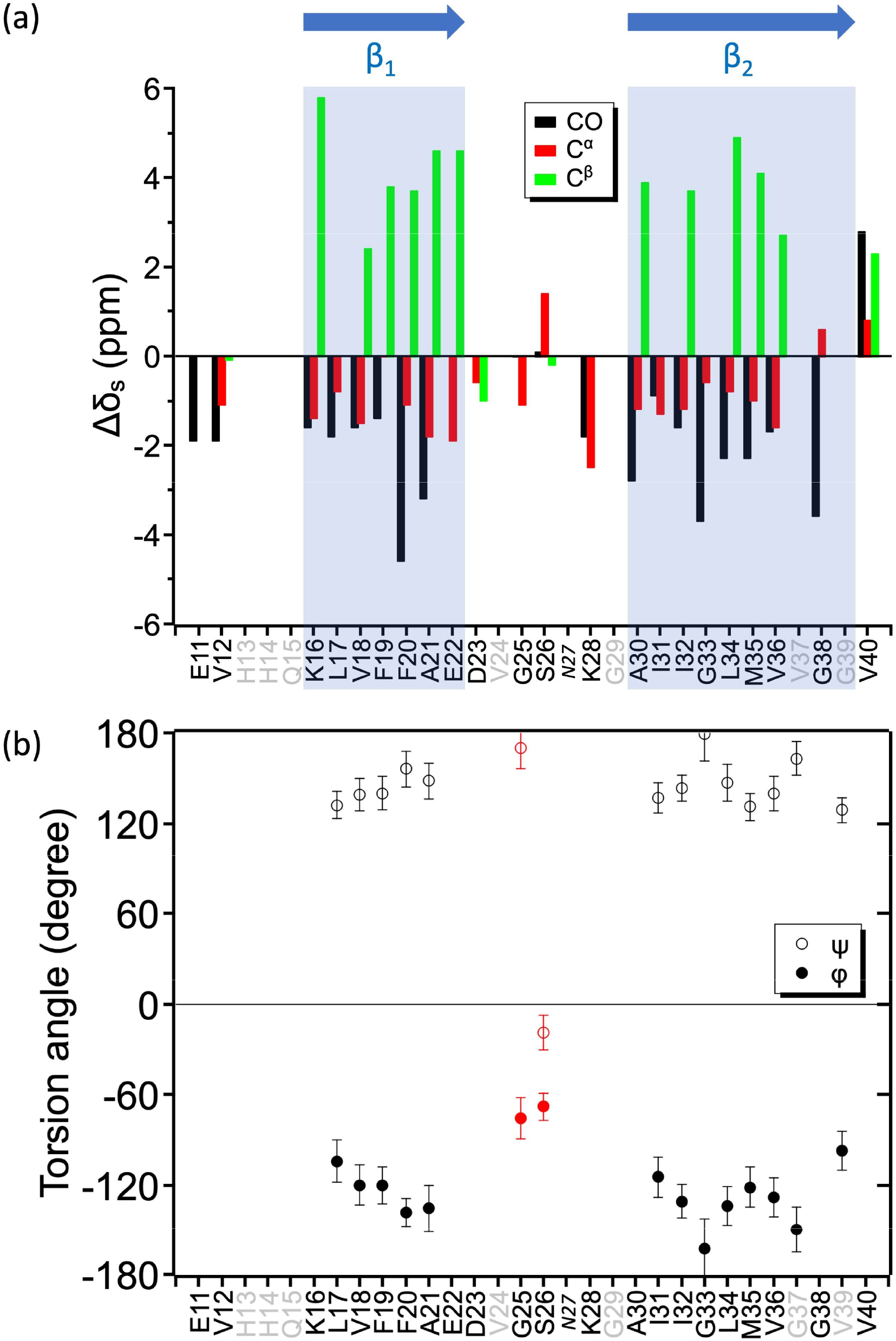
(**a**) Secondary chemical shift (Δ *δ*) of RM10-Aβ_40_. Δ *δ* = *δ* − *δ*, where *δ* denotes the random coiled values. The β-sheet region for RM10-Aβ_40_ was defined under the conditions that Δ *δ* (CO) < 0, Δ *δ* (C) < 0 and Δ *δ* C > 0 were satisfied. The β_1_ and β_2_ regions of RM10-Aβ_40_ were assigned to the residues of K16–E22 and A30–V36, respectively. (**b**) Backbone torsion angles predicted by TALOS-N._[1]_ The data in red belonged to the class of “WARN”, other were “STRONG”.

**Fig. S12.**
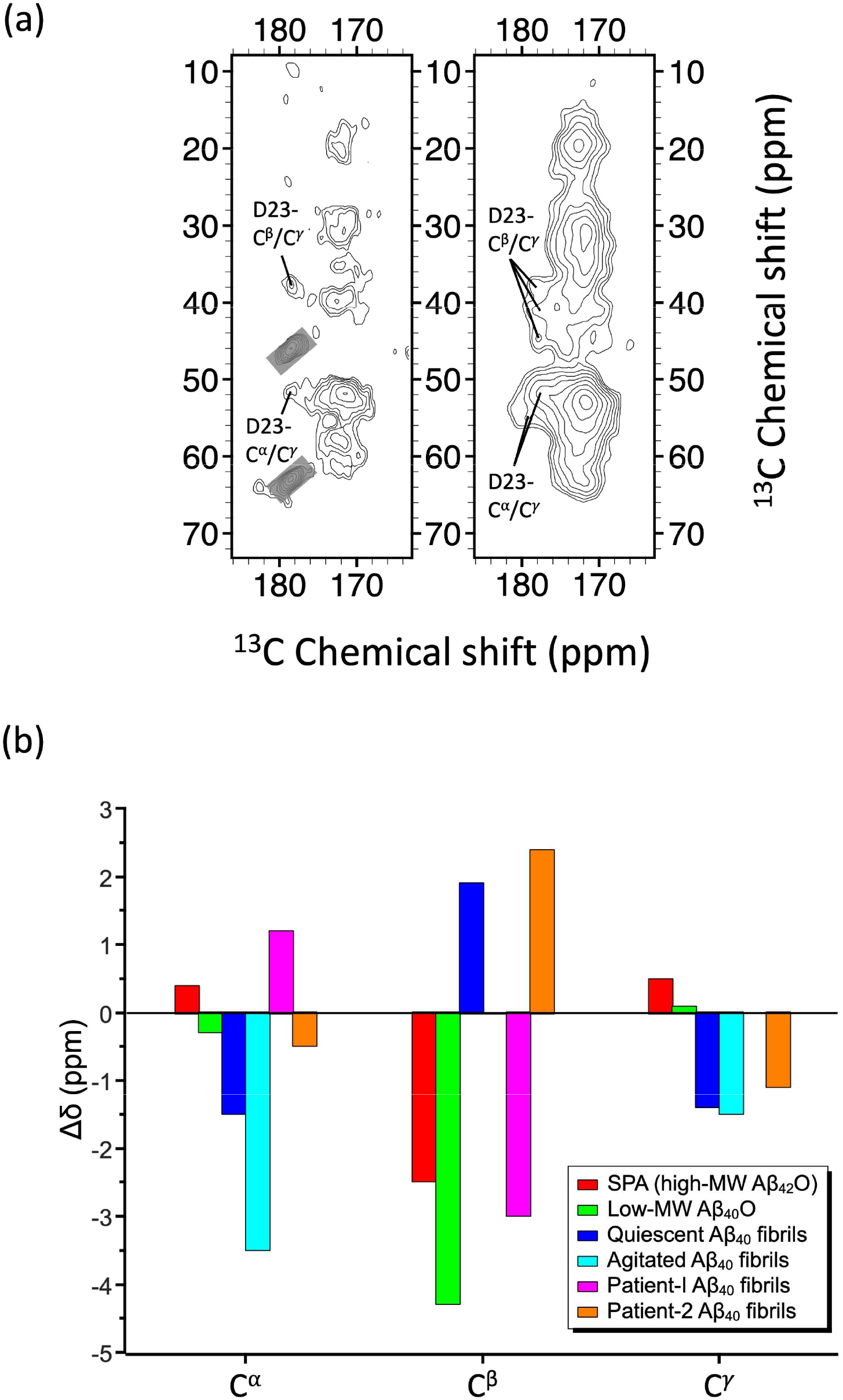
(**a**) Comparison of D23 cross peaks of RM10-Aβ_40_ (left) and RM23-Aβ_40_ (right). The signals masked by the rectangular boxes were spinning sidebands. (**b**) The chemical shifts determined for D23 of RM10-Aβ_40_ were different from the reported data for D23 of other Aβ_40_ aggregates, viz., wild-type Aβ_40_ fibrils incubated under quiescent and agitated conditions,^[2]^ and brain-derived Aβ_40_ fibrils, viz., patient-I,^[3]^ and patient-2,^[4]^ low-MW Aβ_40_Os,^[5]^ and SPA.^[6]^ Note that the D23-CO chemical shift was not available for RM10-Aβ_40_.

**Fig. S13.**
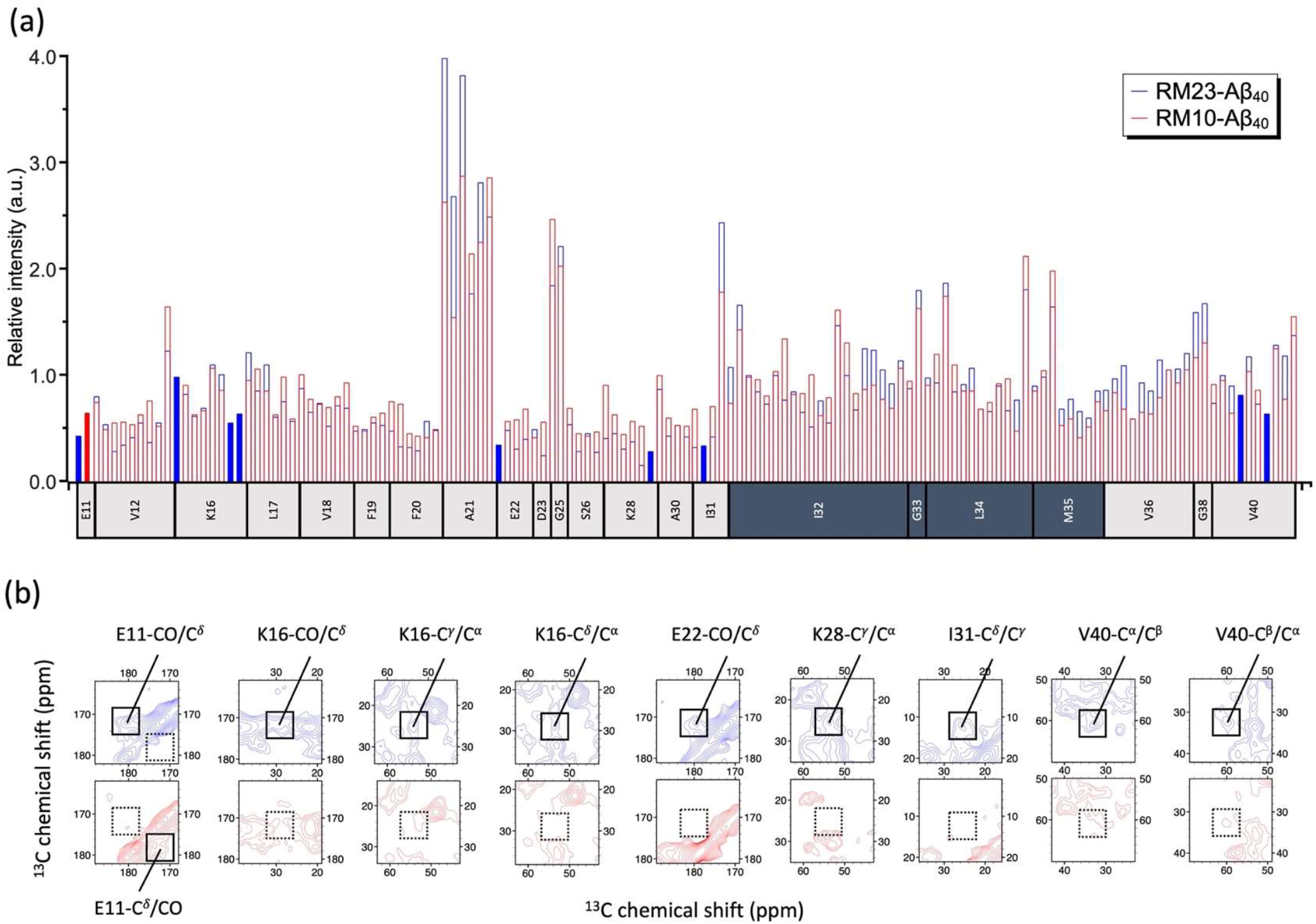
(a) Summary of all the cross-peak intensities for RM23-Aβ_40_ (blue) and RM10-Aβ_40_ (red). The intensities of the two spectra were scaled by minimizing their difference with respect to I32, G33, L34, and M35 (residues in dark grey). Solid blue bars denote the cross peaks that appeared only in the RM23-Aβ_40_ spectrum and the red bar for RM10-Aβ_40_ spectrum. (b) Excerpt spectra of RM23-Aβ_40_ (blue) and RM10-Aβ_40_ (red). The solid and dotted squares indicated the same spectral region, where a cross peak was present and absent, respectively.

**Fig. S14.**
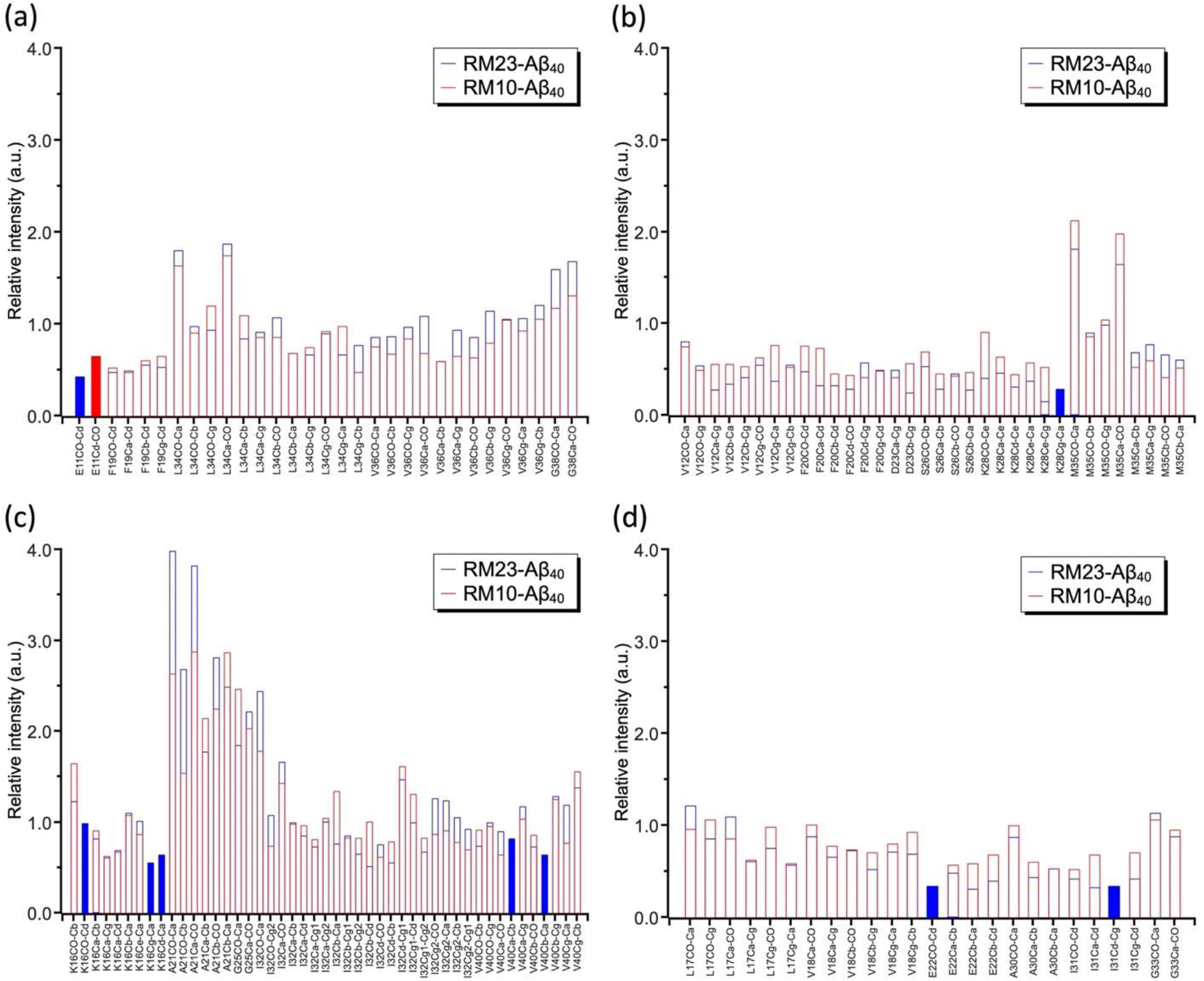
Detailed assignments of the scaled cross-peak intensities of RM23-Aβ_40_ (blue) and RM10-Aβ_40_ (red) shown in Fig. **S13**: (**a**) The intensities corresponding to the S1 sample were scaled by the signal intensities at L34; (**b**) the intensities of the S2 sample were scaled by M35; (**c**) the S3 sample were scaled by I32; (**d**) the S4 sample by G33. Solid bars in blue or red denote the cross peaks that were only observed in the RM23-Aβ_40_ or RM10-Aβ_40_ spectra, respectively.

## Tables

**Table S1.**
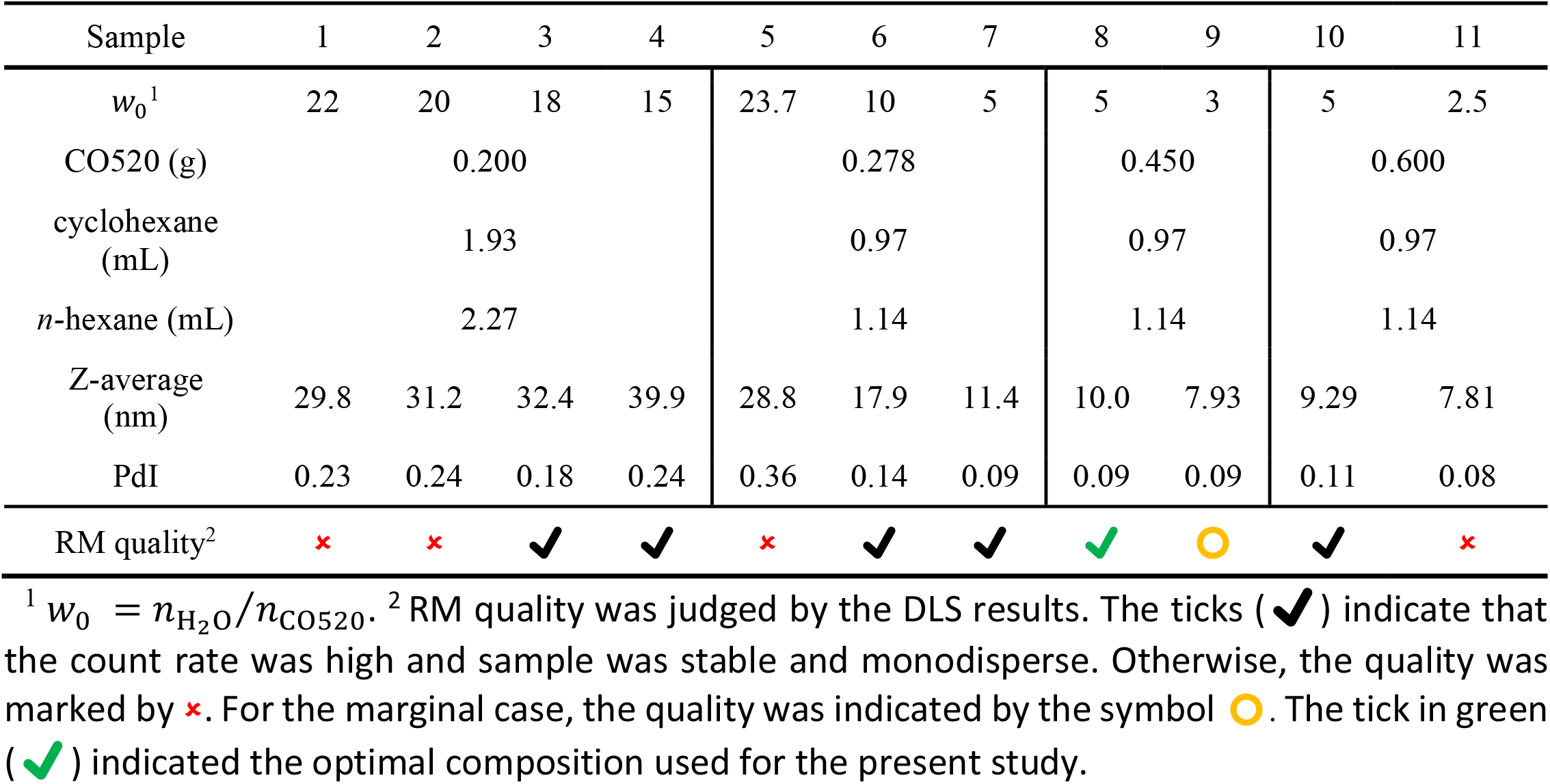
Optimization of the composition of RMs.

**Table S2.**
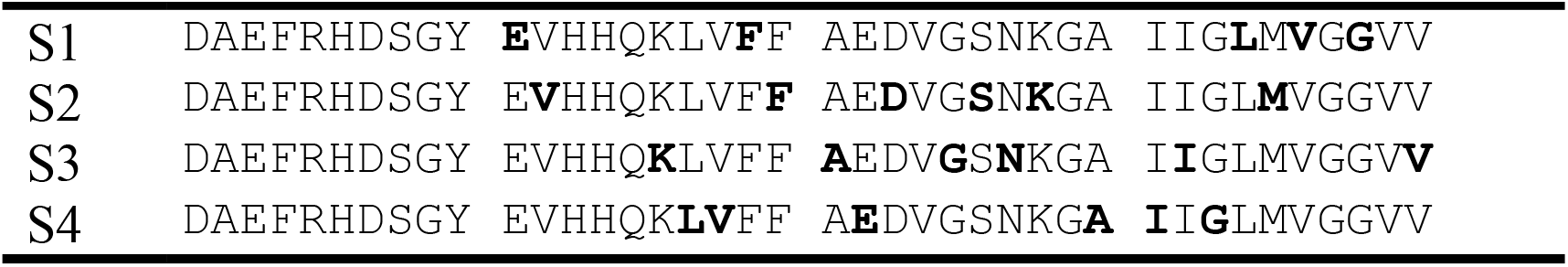
Labeling scheme for the ^13^C and ^15^N enriched RM10-Aβ_40_ sample.

**Table S3.** NMR acquisition parameters used for NMR experiments of the RM10-Aβ_40_.

| Sample | S1 | S2 | S3 | S4 |
| --- | --- | --- | --- | --- |
| B<sub>0</sub> (T) | 14.1 | 14.1 | 9.4 | 9.4 |
| Rotor diameter (mm) | 1.9 | 2.5 | 2.5 | 2.5 |
| $\nu_{\text{MAS}}$ (kHz) | 25.0 | 20.0 | 20.0 | 20.0 |
| Temperature (K) | 268 | 268 | 268 | 268 |
| CP duration (ms) | 2.00 | 2.00 | 2.00 | 2.00 |
| $t_1$ sweep width (kHz) | 30.00 | 37.50 | 25.00 | 25.00 |
| Max $t_1$ (ms) | 3.20 | 3.41 | 2.56 | 2.56 |
| DARR mixing (ms) | 200.00 | 200.00 | 200.00 | 200.00 |
| $t_2$ sweep width (kHz) | 100.00 | 100.00 | 100.00 | 100.00 |
| Max $t_2$ (ms) | 20.48 | 10.24 | 20.48 | 20.48 |
| Recycle delay (s) | 1.00 | 1.00 | 2.00 | 2.00 |
| Experimental time (h) | 54.85 | 70.07 | 73.16 | 91.55 |

**Table S4.** ^13^C NMR chemical shifts for RM10-Aβ_40_ and the backbone dihedral angles predicted by TALOS-N.

| Residue | Chemical shift (ppm)<sup>1</sup> | | | | | | Predicted $\phi, \psi$ (°)<sup>2</sup> |
| --- | --- | --- | --- | --- | --- | --- | --- |
| | CO | C$^{\alpha}$ | C$^{\beta}$ | C$^{\gamma}$ | C$^{\delta}$ | C$^{\epsilon}$ | |
| E11 | 172.2 | nd<sup>3</sup> | nd | nd | 178.8 |  |  |
| V12 | 171.8 | 58.7 | 30.6 | 19.5 |  |  |  |
| K16 | 172.5 | 53.0 | 36.4 | 23.6 | 31.4 | 39.9 |  |
| L17 | 172.8 | 52.0 | nd | 24.5 |  |  | -104 ± 14, 132 ± 9 |
| V18 | 172.2 | 58.3 | 33.1 | 19.8 |  |  | -120 ± 13, 139 ± 11 |
| F19 | 171.9 | 54.9 | 41.2 | 137.0 | 129.6 |  | -120 ± 12, 140 ± 11 |
| F20 | 168.7 | 53.8 | 41.1 | 136.0 | 128.5 |  | -138 ± 10, 156 ± 12 |
| A21 | 172.2 | 48.8 | 21.7 |  |  |  | -136 ± 15, 148 ± 12 |
| E22 | nd | 52.4 | 32.6 | nd | 180.5 |  |  |
| D23 | nd | 51.5 | 37.8 | 178.4 |  |  |  |
| V24 |  |  |  |  |  |  |  |
| G25 | 172.3 | 42.2 |  |  |  |  | -76 ± 14, 170 ± 14 |
| S26 | 172.4 | 57.7 | 61.7 |  |  |  | -68 ± 9, -19 ± 11 |
| N27 | nd | nd | nd | nd |  |  |  |
| K28 | 172.3 | 51.9 | nd | 24.7 | 28.5 | 40.0 |  |
| G29 |  |  |  |  |  |  |  |
| A30 | 172.6 | 49.5 | 20.9 |  |  |  |  |
| I31 | 172.6 | 57.3 | nd | 26.2, nd | 12.9 |  | -100 ± 12, 132 ± 9 |
| I32 | 171.9 | 57.4 | 39.9 | 25.9, 15.9 | 12.7 |  | -128 ± 9, 141 ± 11 |
| G33 | 168.6 | 42.8 |  |  |  |  | -158 ± 23, 174 ± 17 |
| L34 | 172.3 | 52.1 | 44.8 | 25.0 |  |  | -129 ± 10, 140 ± 12 |
| M35 | 171.6 | 52.1 | 35.0 | 33.5 |  |  | -126 ± 11, 135 ± 10 |
| V36 | 172.1 | 58.2 | 33.4 | 19.5 |  |  | -130 ± 13, 139 ± 11 |
| G37 |  |  |  |  |  |  | -150 ± 15, 163 ± 11 |
| G38 | 168.7 | 44.0 |  |  |  |  |  |
| V39 |  |  |  |  |  |  | -98 ± 14, 132 ± 10 |
| V40 | 176.5 | 60.6 | 33.0 | 20.0 |  |  |  |

<sup>1</sup> Chemical shifts referenced to neat TMS. <sup>2</sup> The data in red belonged to the class of “WARN”, other were “STRONG”. <sup>3</sup> nd: not determined.

## Notes

Supporting information for this article is given via a link at the end of the document.

### Competing Interest Statement

The authors have declared no competing interest.

